# Ecdysone-mediated intestinal growth contributes to microbiota-driven developmental plasticity under malnutrition

**DOI:** 10.1101/2025.02.07.637066

**Authors:** Longwei Bai, Stéphanie Bellemin, Elodie Guillemot, Maura Strigini, Benjamin Gillet, Cathy Isaura Ramos, François Leulier

**Affiliations:** Institut de Génomique Fonctionnelle de Lyon, Ecole Normale Supérieur de Lyon, CNRS UMR 5242, Université Claude Bernard Lyon 1, 46 allée d’Italie, 69007 Lyon, France; Synbird, 14 Fbg Reclus, 73000 Chambéry, France; Charles River Laboratories, 9 All. Moulin Berger, 69130 Écully, France; Université Jean Monnet Saint-Étienne, Mines Saint Etienne, INSERM, SAINBIOSE U1059, 42023 Saint-etienne, France

**Keywords:** Adaptive growth, developmental plasticity, ecdysone signaling, microbiota, intestine, *Drosophila*

## Abstract

Organ and systemic growth must remain coordinated during development, even under nutritional stress. In *Drosophila* larvae, the intestinal microbiota contributes to this coordination by promoting growth and maturation under chronic undernutrition. Using gnotobiotic models, we show that association with *Lactiplantibacillus plantarum* (*Lp*) selectively enhances midgut growth relatively to other organs, providing an adaptive mechanism that buffers the impact of dietary restriction. Transcriptomic profiling of larval midguts revealed a strong Ecdysone signaling signature upon *Lp* association. Functional analyses showed that local conversion of Ecdysone to its active form, 20-hydroxyecdysone, by the cytochrome P450 enzyme Shade, together with enterocyte Ecd receptor activity, is required for *Lp*-dependent intestinal and systemic growth. Pharmacological activation of Ecd signaling partially mimicked the bacterial effect, confirming its sufficiency to drive adaptive midgut expansion. Our results uncover an unexpected role of intestinal Ecd signaling in microbiota-driven developmental plasticity, revealing how commensal bacteria modulate local steroid signaling to fine-tune organismal growth and maturation.

## Introduction

In animals, organ growth is coordinated with systemic growth to ensure synchronized development. Beyond this coordination, organisms exhibit developmental plasticity, the ability to adjust growth and maturation rates in response to environmental and internal cues, ensuring the emergence of fit adults (Pigliucci, 2001). Nutrition is a major regulator of developmental plasticity, and the microbiota has emerged as another key determinant of this process (Storelli et al., 2011, Shin et al., 2011).

In *Drosophila* larvae, nutrient restriction delays growth and development. The absence of a microbiota further exacerbates this delay, whereas mono-association of germ-free (GF) animals with *Lactiplantibacillus plantarum* (*Lp*), a representative member of the fly microbiota, restores both growth and maturation rates (Storelli et al., 2011). These beneficial effects of *Lp* depend on enhanced digestive capacity and improved systemic growth signaling under chronic undernutrition (Erkosar et al., 2015, Matos et al., 2017, Storelli et al., 2018).

Adaptive developmental responses in *Drosophila* are primarily governed by two hormonal systems: Insulin-like peptides (Ilps), which control systemic growth rate (Colombani et al., 2003, Geminard et al., 2009), and the steroid hormone ecdysone (Ecd), which determines the duration of growth periods by triggering molting and maturation (Shingleton et al., 2005). Microbiota-dependent developmental acceleration involves both axes through TORC1 activation in the fat body and prothoracic gland (PG) (Storelli et al., 2011).

Ecd is synthesized from dietary cholesterol in the PG under control of the prothoracicotropic hormone, secreted into the hemolymph, and taken up by peripheral tissues through Organic Anion Transporting Polypeptides such as the Ecdysone Importer EcI/Oatp74D (Okamoto et al., 2018, Okamoto and Yamanaka, 2020). Once internalized, Ecd is converted by the cytochrome P450 enzyme Shade into its active form, 20-hydroxyecdysone (20E), which binds to the Ecdysone receptor (EcR) to regulate gene expression (Petryk et al., 2003, Yamanaka et al., 2015).

While the systemic and local actions of Ilps are well understood, the tissue-specific contributions of Ecd remain poorly characterized, owing to its pleiotropic effects on growth, remodeling, and differentiation across larval and imaginal tissues (Boulan et al., 2015). How these canonical systemic signals translate into tissue-specific growth responses during nutritional stress remains largely unknown. Here, we identify intestinal Ecd signaling as a key mediator of microbiota-dependent developmental plasticity. Transcriptomic and functional analyses reveal that *Lp* association enhances Ecd signaling in larval enterocytes during malnutrition, promoting adaptive intestinal growth that contributes to systemic developmental coordination.

## Results and Discussion

### Microbiota association selectively promotes midgut growth under malnutrition

To assess how *Lp* influences organ growth during chronic malnutrition, we compared the growth of diploid (central nervous system, wing discs) and polyploid (salivary glands, midgut) larval organs in GF and *Lp*-associated conditions (Fig. 1A). While the CNS, wing discs, and salivary glands were comparable between GF and *Lp*-associated larvae (Fig. 1B–D), the midgut displayed a robust increase in length (Fig. 1E–F). Thus, the larval midgut is a primary target of microbiota-mediated growth under nutrient restriction.

**Figure 1.**
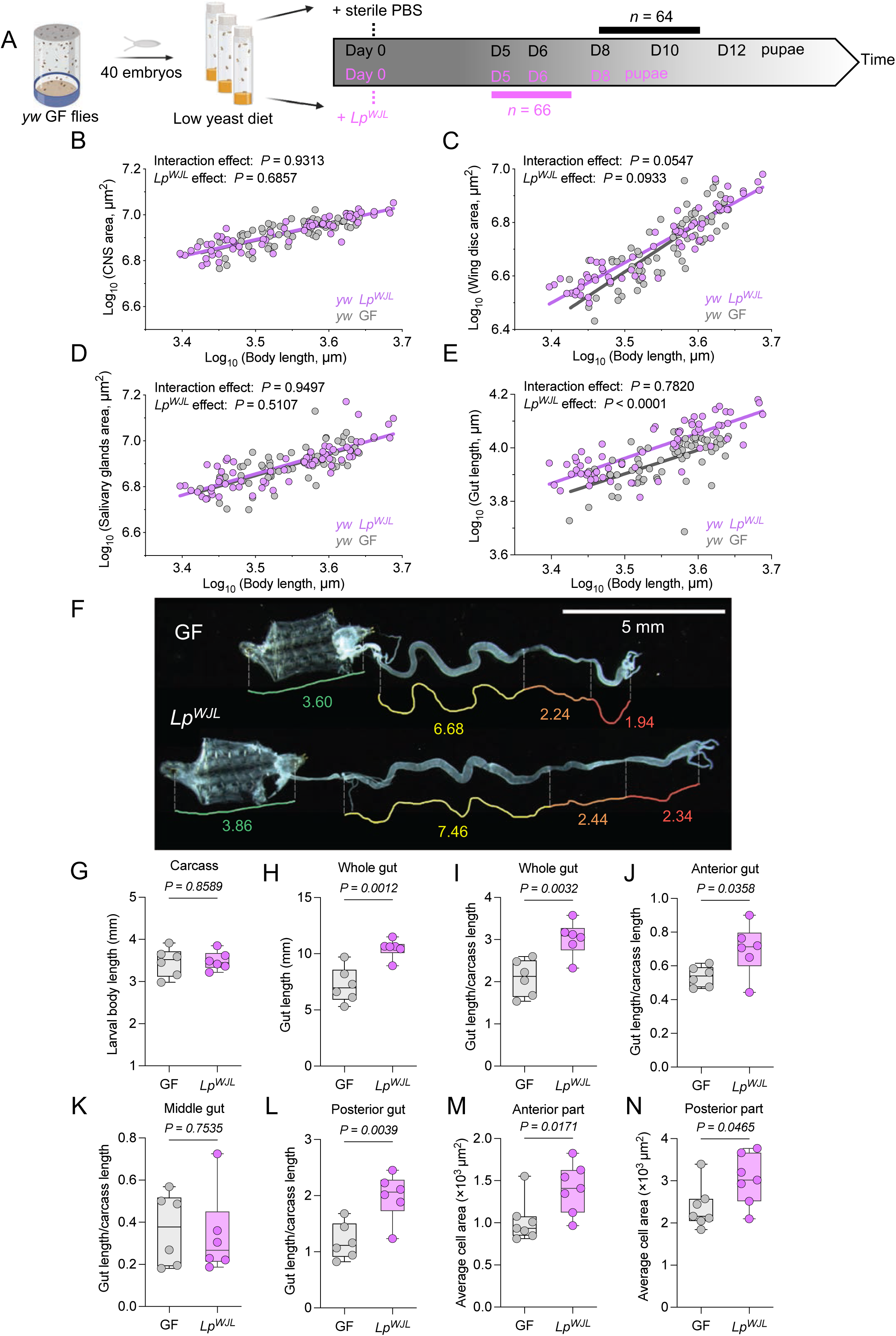
Microbiota association selectively promotes midgut growth under malnutrition. (A) Experimental roadmap detailing organ collection for L3 instar *yw* larvae: *Lp^WJL^*-associated (D5–D7) and GF (D8–D11). *n* = number of dissected larvae. (B–E) Allometric scaling of organ size/organ area vs. body length for: (B) CNS, (C) imaginal wing discs, (D) salivary glands, and (E) midguts. Simple linear regressions are shown for each group. *P* values for the allometric interaction (slope difference) and the *Lp^WJL^* effect (elevation difference) are indicated. (F) Representative images of dissected guts attached to their respective carcasses (body length readout, green line) from *Lp^WJL^*-associated (D6) and GF larvae (D9). Red, orange, and yellow lines indicate the anterior, middle, and posterior gut regions, respectively. Scale bar represents 5 µm. (G–L) Quantification of gut size in size-matched GF and *Lp^WJL^*-associated larvae. (G) Larval size. (H) Whole gut length. (I–L) Normalized gut length (gut length/carcass length ratio) for the whole gut (I), anterior gut (J), middle gut (K), and posterior gut (L). *n* = 6. (M–N) Quantification of enterocyte cell area in the anterior (M) and posterior (N) gut regions. *n* = 7. In all panels, purple indicates *Lp^WJL^*-associated larvae and GF larvae in gray. Statistical significance was determined using a two-tailed unpaired *t*-test (G–J, L, N) and a lognormal *t*-test (K, M). *P* values are indicated on the panels.

To determine whether this growth occurred uniformly along the gut, we examined anterior, middle, and posterior regions separately. Despite *Lp* colonizing mainly the anterior half of the midgut (Storelli et al., 2018), both anterior and posterior regions were significantly longer in *Lp*-associated larvae (Fig. 1G–L). Increased enterocyte size in these regions (Fig. 1M–N; Fig. S1) indicates that *Lp*-mediated midgut growth results at least in part from enhanced cell growth.

These findings demonstrate that *Lp* association triggers a coordinated growth response within the larval midgut. The spatial distribution of this effect, which does not mirror bacterial abundance, suggests that *Lp*-induced signals may propagate along the gut or act through systemic factors. Recent work in the adult intestine (Blackie et al., 2024) highlights functional coupling between distinct epithelial compartments through spatial proximity and signaling cross-talk, raising the possibility that folded larval midgut regions also interact dynamically within the larval milieu. Furthermore, the observation that both anterior and posterior but not the middle segment enlarge upon *Lp* association suggests potential functional heterogeneity among intestinal cell populations, possibly linked to differential hormonal responsiveness.

### Microbiota association induces an intestinal ecdysone signaling signature

To uncover transcriptional programs underlying microbiota-dependent intestinal growth, we performed RNA sequencing on midguts dissected from size-matched GF and *Lp*-associated larvae at early and late third-instar (L3) stages (Fig. 2A, Fig. S2A–B). This design controlled for *Lp*-induced developmental acceleration. Transcriptomic profiling identified 2,633 differentially expressed genes across both stages. Functional clustering (DAVID) yielded 91 annotation clusters, with the strongest enrichment for “Ecdysteroid kinase-like” genes, followed by categories related to mitochondrial activity, proteolysis, and signal transduction (Fig. S2C–D). Gene Ontology analysis revealed a unique enrichment for the “Response to ecdysone” term (Fig. S3), and several canonical ecdysone-responsive genes were upregulated in *Lp*-associated midguts (Fig. S4A).

**Figure 2.**
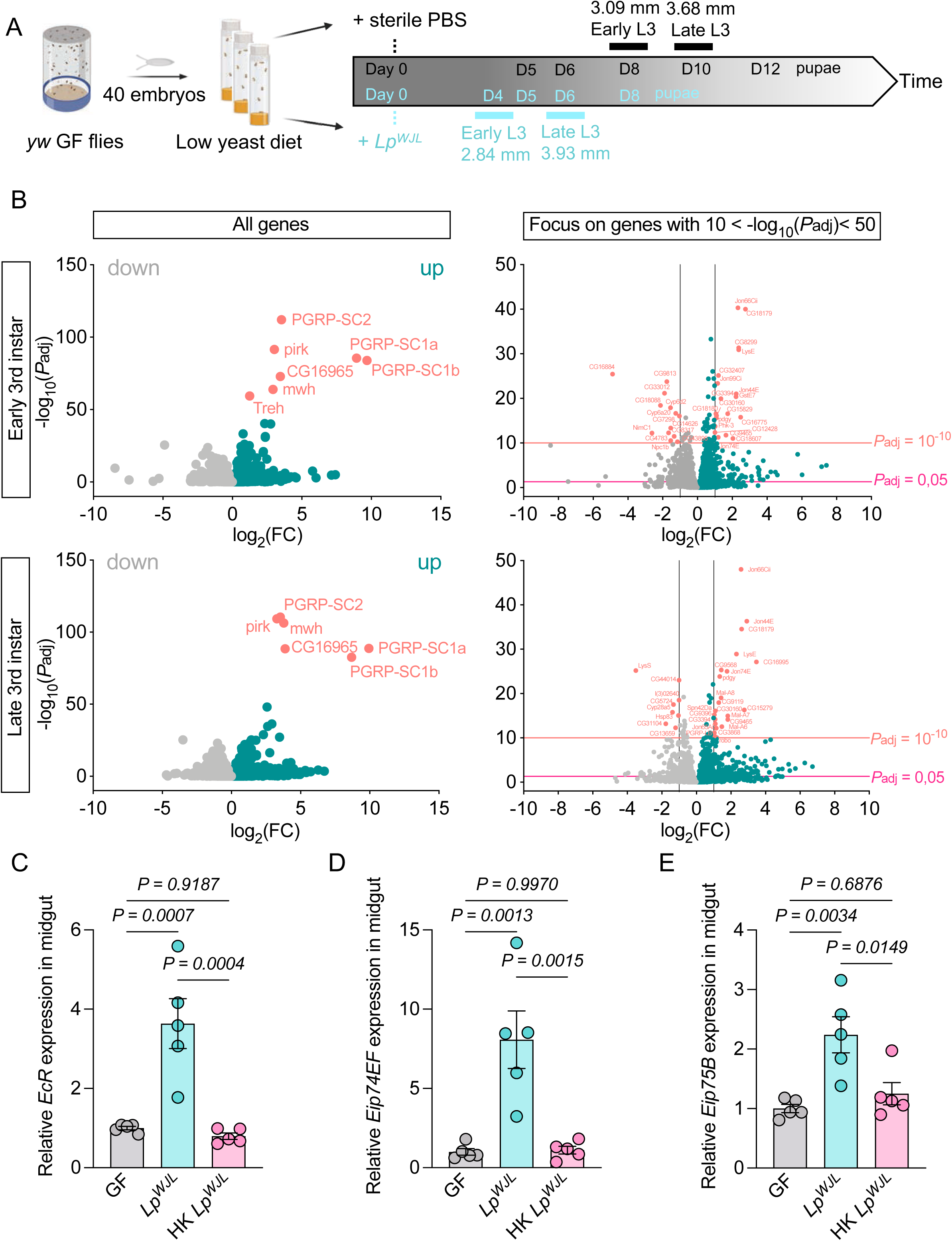
Microbiota association induces an intestinal ecdysone signaling signature. (A) Experimental roadmap detailing midgut collection from size-matched *Lp^WJL^*-associated and GF larvae at two developmental stages: early L3 and late L3. (B) Volcano plots illustrating differential gene expression between *Lp^WJL^*-associated and GF midguts at early L3 (upper panel) and late L3 (bottom panel) instars. All genes are shown in the left panel, while the right panel highlights genes with a strong significance (10 < -log10 (*P*_adj_) < 50). (C–E) Relative transcript levels of ecdysone-responsive genes in the gut—*EcR* (C), *Eip74EF* (D), and *Eip75B* (E) were analyzed by qRT-PCR in GF, *Lp^WJL^*-associated, and heat-killed (HK) *Lp^WJL^*-associated larvae under malnutrition. Data are presented as mean ± SEM. Statistical significance was determined using one-way ANOVA with Tukey’s multiple comparisons test. *P* values are indicated on the panels. *N* = 5.

Differential expression was extensive at both stages (early L3: 1,624 genes; late L3: 1,462 genes), with a predominance of upregulated transcripts (Fig. 2B). As expected, *Lp* association induced IMD pathway targets (*PGRP-SC1*, *PGRP-SC2*, *Pirk*) and markers of intestinal maturation (*jon66cii*, *jon44E*; Fig. S5A–B). In contrast, canonical growth regulators (*Dilps*, *dMyc*, *Hippo*, *EGFR*) were unchanged (Fig. S4B), indicating that microbiota-driven midgut growth operates through alternative mechanisms. RT-qPCR confirmed the RNA-seq results, showing increased expression of EcR target genes in midguts from *Lp*-associated larvae but not from animals colonized with heat-killed bacteria (Fig. 2C–E). These findings demonstrate that live *Lp* bacteria are required to induce intestinal Ecd signaling under malnutrition. Altogether, these data reveal that *Lp* association activates a distinct intestinal ecdysone signaling program, identifying Ecd as a candidate hormonal mediator of microbiota-driven adaptive growth.

### Enterocyte ecdysone signaling mediates microbiota-dependent intestinal growth

Because *Lp* promotes systemic growth and induces intestinal Ecd targets, we next examined whether Ecd signaling is required for midgut adaptation. Quantification of 20E showed significant elevated levels in both midgut and hemolymph of size-matched *Lp*-associated larvae compared with GF controls, whereas fat body levels were slightly elevated (Fig. 3A–C).

**Figure 3.**
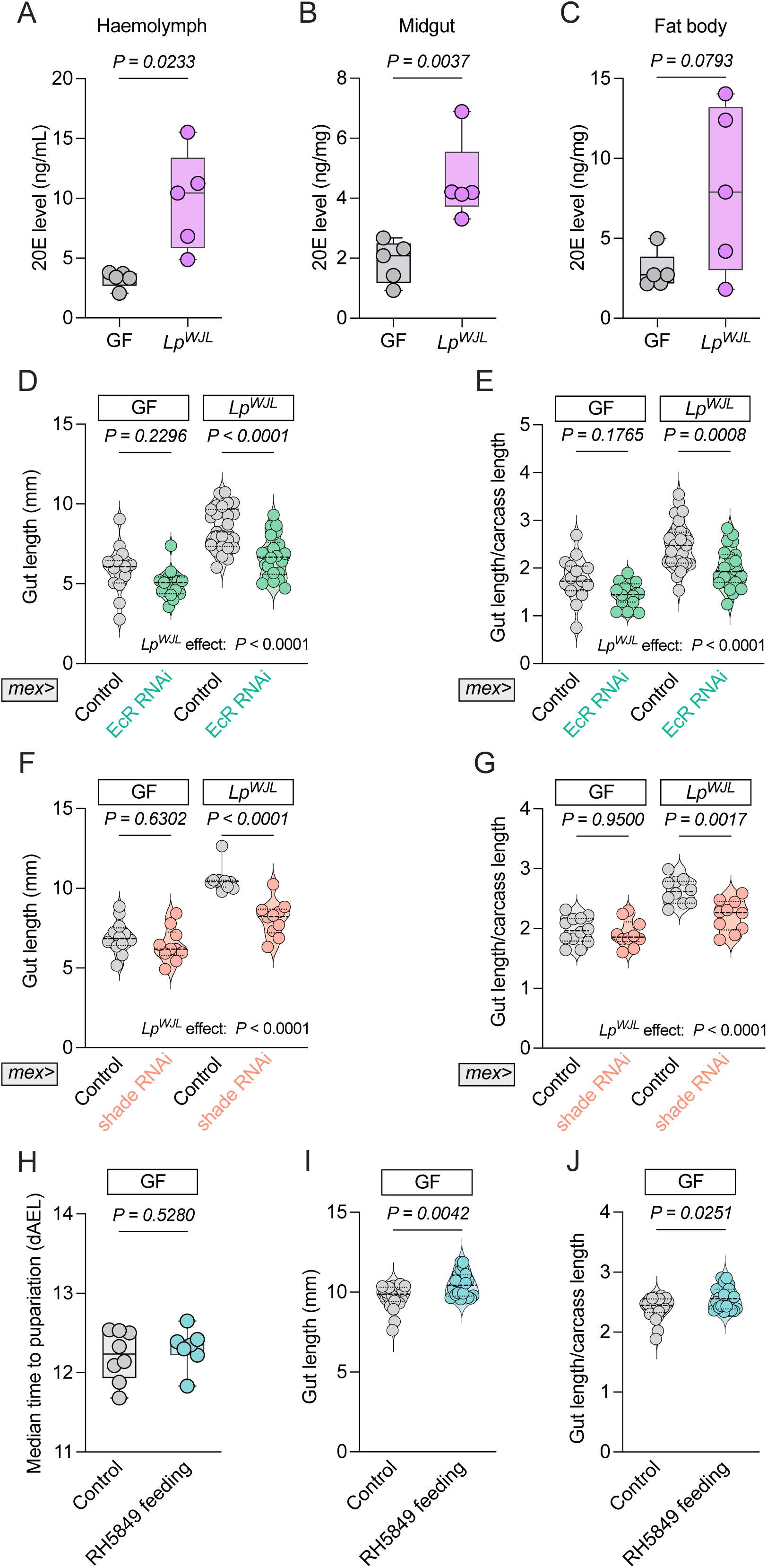
Enterocyte ecdysone signaling mediates microbiota-dependent intestinal growth. (A–C) Quantified 20E levels in the haemolymph (A), midgut (B), and fat body (C) of size-matched GF and *Lp^WJL^*-associated larvae. *n* = 5. (D–G) Measured gut length in size-matched GF and *Lp^WJL^*-associated larvae with *EcR* knockdown in enterocytes (ECs) (*mex>EcR* RNAi) (D) and *shade* knockdown (*mex>shade* RNAi) (F). Normalized gut length with *EcR* RNAi (E) and *shade* RNAi (G) was also determined. Controls included respective TRiP and KK lines. *n* ≥ 10. (H–J) Quantified median time to pupariation (H), gut length (I), and normalized gut length (J) in *yw* GF larvae fed the ecdysone agonist RH5849 compared to a MeOH vehicle control. *n* ≥ 7. Statistical significance was determined using a two-tailed Welch’s *t* test (A), a lognormal *t*-test (B–C), a two-way ANOVA with Tukey’s multiple comparisons test (D–G), a two-tailed unpaired *t* test (H), and a two-tailed Mann-Whitney *U* test (I–J). *P* values are indicated on the panels.

Reducing *EcR* expression specifically in enterocytes by RNAi markedly suppressed *Lp*-dependent midgut growth and gut-to-carcass ratios (Fig. 3D–E; Fig. S5C), demonstrating that enterocyte EcR activity is necessary for the bacterial effect. Similarly, silencing *shade* also reduced *Lp*-dependent growth (Fig. 3F–G; Fig. S5D). Because Shade is enriched in the larval midgut (Chung et al., 2009, Brown et al., 2014), these results indicate that *Lp* enhances local Ecd activation in enterocytes.

Finally, dietary supplementation with the nonsteroidal Ecd agonist RH5849 (Wing, 1988) modestly increased midgut length in GF larvae without affecting developmental timing (Fig. 3H–J), indicating that activation of intestinal Ecd signaling is sufficient to partially reproduce the *Lp*-induced intestinal growth effect.

### Intestinal ecdysone signaling supports systemic growth adaptation during microbiota association

To test whether intestinal Ecd signaling also contributes to systemic developmental outcomes, we silenced *EcR* or *shade* in specific tissues and measured developmental timing and pupal size during malnutrition (Fig. 4; Fig. S6A–D). Disrupting Ecd signaling in the fat body using *Lpp*-GAL4 slightly delayed development in both GF and *Lp*-associated larvae and triggered full pupal lethality in all conditions, stressing the essential role of fat-body Ecd responses for proper developmental progression and metamorphosis. In contrast, enterocyte-specific *EcR* knockdown delayed pupariation only in GF animals (Fig. 4A,C), while *Lp*-associated counterparts developed as controls. *shade* knockdown did not affect timing in either condition (Fig. 4B,D). Notably, both *EcR*- and *shade*-RNAi *Lp*-associated larvae produced significantly smaller pupae than controls (Fig. 4E–F).

**Figure 4.**
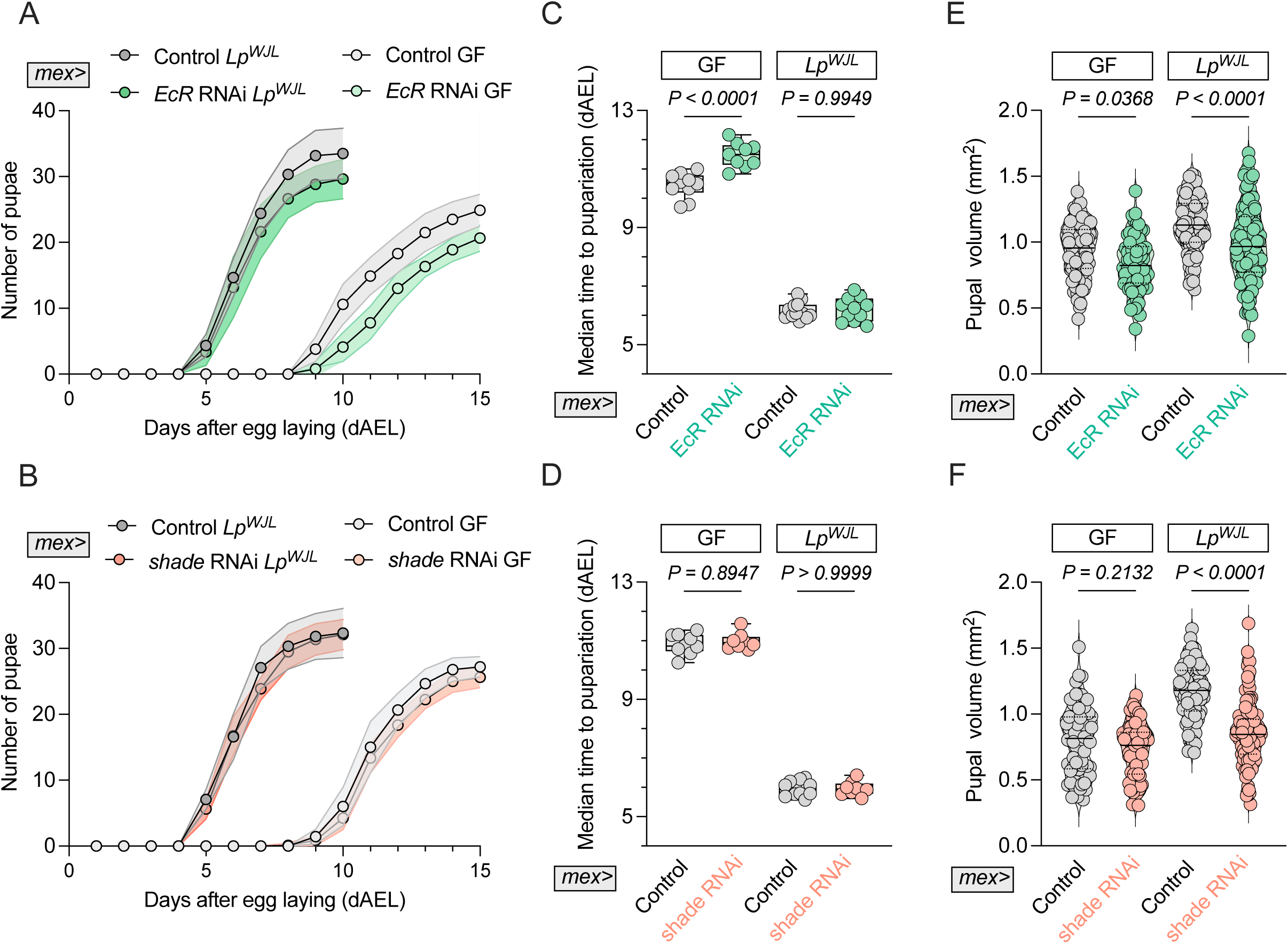
Intestinal ecdysone signaling supports systemic growth adaptation during microbiota association. Under malnutrition, measured developmental timing (A–B), corresponding median time to pupariation (C–D), and pupal volumes (E–F) are shown in GF and *Lp^WJL^*-associated larvae with *EcR* knockdown in ECs (*mex>EcR* RNAi) (A, C, E) and *shade* knockdown (*mex>shade* RNAi) (B, D, F). Controls included respective TRiP and KK lines. Statistical significance was determined using a two-way ANOVA with Tukey’s multiple comparisons test. *P* values are indicated on the panels. (A–D): *n* ≥ 8. (E–F): *n* ≥ 65.

These findings indicate that while fat-body Ecd signaling regulates developmental tempo and ensures developmental transitions such as metamorphosis, intestinal Ecd activity specifically supports systemic growth adaptation during *Lp* association. Thus, microbiota-enhanced enterocyte Ecd signaling contributes to both local gut expansion and organismal growth plasticity under nutrient restriction.

### Conclusion

Together, our findings reveal that organ and systemic growth remain tightly coordinated even under nutritional stress, and that this coordination relies on microbiota-mediated modulation of intestinal physiology. Under malnutrition, association with *Lactiplantibacillus plantarum* promotes enterocyte growth and intestinal expansion, providing an adaptive mechanism that buffers the impact of dietary restriction on developmental growth.

Transcriptomic and functional analyses identify intestinal Ecd signaling as a key mediator of this adaptive response. *Lp* association enhances the local conversion of Ecd to its active form 20E, through Shade activity, and both Shade and the EcR are required for *Lp*-dependent intestinal and systemic growth. These results establish intestinal Ecd signaling as a central hormonal component of microbiota-driven developmental plasticity.

Recent studies have emphasized that Ecdysone signaling in adult enterocytes orchestrates cellular hypertrophy and endoreplication to control midgut size and functionality (Ahmed et al., 2020, Zipper et al., 2025, Zipper et al., 2020). Integrating these findings with our work suggests that the microbiota leverages this local Ecdysone-dependent growth program to tune intestinal capacity according to nutritional availability in the context of larval development. This highlights Ecdysone signaling in enterocytes as a pivotal interface between microbial cues, tissue plasticity, and systemic developmental control and calls for investigating the contribution of microbiota to adult intestinal plasticity.

By uncovering a direct link between commensal bacteria, intestinal steroid signaling, and systemic growth control, this work expands the physiological framework of Ecd action beyond its classical endocrine role. It highlights how local steroid signaling within an epithelial organ can exert systemic influence on growth, revealing a new level of developmental coordination between tissues and their microbial partners. Similar microbiota–steroid interactions occur in mammals, where gut bacteria, including *Lactobacillus* species, produce β-glucuronidase enzymes that reactivate estrogens (Baker et al., 2017, Ervin et al., 2019, Hu et al., 2023). Given the functional analogies between Drosophila Ecdysone and mammalian estrogens (Aranda and Pascual, 2001, Martinez et al., 1991, Oberdorster et al., 2001), our findings suggest that microbiota-mediated modulation of steroid activity represents an evolutionarily conserved mechanism linking microbial cues to host endocrine control. It positions the larval midgut as an adaptive interface integrating nutritional, microbial, and hormonal cues to fine-tune organismal maturation.

## Methods

### Fly stocks and media

Conventional *Drosophila* stocks were reared at 25°C under a 12:12 hour light/dark cycle using a standard cornmeal-yeast medium containing 8.2 g/L agar (#20768.361, VWR), 80 g/L cornmeal flour (Westhove, Farigel maize H1), 80 g/L inactivated yeast (#24979.413, VWR), and supplemented with 5.2 g/L Methyl 4-hydroxybenzoate sodium salt (#106756, Merck) and 4 mL/L of propionic acid (#409553, Carlo Elba). The axenic embryos were generated as previously described (Ma et al., 2015). These sterilized embryos were subsequently raised on a standard medium that included a cocktail of four antibiotics (Ampicillin, Kanamycin, Tetracycline, and Erythromycin) to maintain a GF environment for the newly hatched larvae. The nutritional challenge was performed using fresh food where the amount of inactivated yeast was reduced to 7 g/L (referred as to low-yeast diet). The following fly stocks were used: *yw* (obtained from Bruno Lemaitre), *mex*-*GAL4* (obtained from Irene Miguel-Aliaga), *Lpp-GAL4* (obtained from Alexandre Djiane), *UAS-EcR* RNAi (BDSC#9327), *UAS-shade* RNAi (VDRC#108911), and respective TRiP control line (BDSC# 35785) and KK control line (VDRC#60101). All experimental crosses were performed at 29°C and females laid eggs on low-yeast food.

### Bacterial culture and larvae mono-association

*Lp^WJL^* referred to *Lp* in this manuscript (Storelli et al., 2011) was cultured overnight in Man, Rogosa and Sharpe (MRS) broth at 37°C without shaking. For inoculation, tubes containing 40–60 GF embryos were seeded with resupened *Lp* to achieve a final dose of 10^8^ CFUs/tube. As a control, bacteria were inactivated by heat using a 95°C water bath treatment for 45 min. For GF controls, an equal volume of sterile PBS was applied.

### RNA sequencing

GF or *Lp*-associated *yw* axenic embryos were reared on a low-yeast diet. Gut collection from L3 larvae was performed to obtain size-matched larvae: early L3 samples (15 guts/replicate, 5 replicates) were collected at 4 days AEL for *Lp* and 8 days AEL for GF, while late L3 samples (10 guts/replicate, 5 replicates) were collected at 6 days AEL for *Lp* and 10 days AEL for GF. RNA was extracted using the NucleoSpin kit (Macherey Nagel, #740955) following the manufacturer’s protocol, and quantified using the Nanodrop spectrophotometer (Thermofisher, ND-2000). For RNA sequencing, samples were treated by the IGFL sequencing platform. Purification, quantification and quality checks were performed using the Qubit 2.0 – RNA BR Assay Kit (Thermofisher) and the TypeStation 2200 (Agilent). Libraries were generated following the online protocol (https://www.hexabiogen.com), quantified via the TapeSation D1000 ScreenTape Assay (Agilent), and the raw data were subsequently processed to generate final genes regulation datasets as in (Grenier et al., 2023). The sequencing data were analyzed using the DAVID Bioinformatics Resources online software (https://david.ncifcrf.gov/) on 2633 annotated genes across both sizes groups, which were organized into a functional annotation chart and subsequently grouped into 91 clusters. An enrichment score (ES) was calculated (ES = -log_10_ mean of the adjusted *P* value (*P*_adj_)) and 53 of the clusters were found to be significantly enriched (ES >1.3, corresponding to *P*_adj_ < 0.05). Then, genes from the two sizes were split into early L3 and late L3 groups and further analyzed by Biological Process Direct with *P*_adj_ < 5.1ξ 10^-2^ and presented as pie charts. Finally, volcano plots were generated using GraphPad Prism 10.

### RT-qPCR

RNA was extracted from ten dissected guts of size-matched larvae after homogenization with glass beads in a Precellys 24 and purified using the NucleoSpin kit (Macherey-Nagel). Reverse transcription of 1μg of RNA was performed with SuperScript II (Invitrogen). Quantitative PCR (qPCR) was run on an iQ5 Real-time PCR Detection System (Biorad) using SYBR GreenER qPCR SuperMix (Invitrogen). Specific and unique amplification was confirmed by melting curve analysis. All expression levels were normalized to two reference genes, *Tubulin* and *RpL32* using delta-delta-Ct method. The following primers (5’–3’) were used for qPCR assays: *tubulin* qF TGTCGCGTGTGAAACACTTC, qR AGCAGGCGTTTCCAATCTG; *RpL32* qF ATGCTAAGCTGTCGCACAAATG, qR GTTCGATCCGTAACCGATGT; *jon66Cii* qF AAACTGACCCCGGTCCAC, qR CCTCCTCAGCCGGATAGC; *EcR* qF GCAAGGGGTTCTTTCGACG, qR CGGCCAGGCACTTTTTCAG; *jon44E* qF ACAGCGCTAACCATGTGCT, qR GGTGTACTGGGCCTCGTG; *shade* qF CCCACCAAAACGTACAGGGA, qR GTAATCCAGCAGCCTCGTCA; *Eip74EF* qF CATAAAGACGGAGCAAAATACGC, qR CCGCTAAGCAGATTGTGGAG; *Eip75B* qF ATCTGCATGTTTGACTCGTCG, qR TCCGCGAAATTGAAGGTGGAG.

### Developmental timing determination and pupal volume analysis

40 or 60 (in RNAi conditions) axenic eggs were inoculated with *Lp* or sterile PBS (control) and incubated at 25°C (experiments using *yw* animals) or 29°C (RNAi experiments). Larval developmental timing was assesses by daily counting the number of new-emerged pupae until all pupae emerged. Median time to pupariation was calculated to evaluate larval developmental timing. Pupae were imaged using a stereomicroscope (M205FA, Leica). Measurements for length (L) and width (W) were taken using Fiji software (Version 1.54r), and the pupal volume was calculated using the formula (4/3) π(L/2) (W/2)^2^.

### Measurement of 20E titers

Ten guts or five fat bodies were dissected from *yw* L3 larvae and pooled in 200 µL of MeOH. Samples were ground with a pestle and vortexed. After adding 300 µL of MeOH, samples were centrifuged for 10 min, 13,000g at 4_°_C. The supernatant was kept in a new tube and the pellet was re-extracted with 500 µL MeOH, vortexed and centrifuged. Finally, the last extraction was performed using 500 µL EtOH. The three collected supernatants were pooled and stored at -20_°_C. For normalization, a duplicate of this experiment was performed and samples were quantify using BCA assay kit (SIGMA). For haemolymph, *yw* L3 larvae were dried on tissue paper and placed on parafilm. The cuticle was gently pricked to collect 2 µL haemolymph from 8 larvae that were placed in 200 µL of MeOH. Then, the samples were treated the same way as described above. Most extraction steps were performed on ice. Evaporation of the samples was done using a SpeedVac and then dissolved in EIA buffer from the kit (A05120, Bertin) and treated following the manufacturer’s instructions.

### Nonsteroidal Ecd agonist treatments

*yw* axenic embryos were transferred to freshly prepared low-yeast fly medium, which was supplemented with the nonsteroidal Ecd agonist RH5849 (5 µg/mL) (DRE-C16813000, DrEhrenstorfer) or an equal volume of MeOH as a vehicle control. The flies were then reared at 25°C for the subsequent developmental timing and gut size measurements.

### Organ morphometric analysis

*Lp*-associated larvae (5–7 days AEL) and GF larvae (8–11 days AEL) were collected during the middle to late L3 instar (pre-wandering). Four organs were dissected from each larva, directly oriented on a poly-Lysine coated slide, and imaged using a stereomicroscope (M205FA, Leica). Organ area/length was measured. Data were analyzed using log-log plots to determine the allometric scaling relationship y = bx^α^ (Huxley, 1924).

### Gut morphometry and cell area analysis

L3 instar larval guts were dissected, immediately fixed for 1 min in Bouin’s solution (HT10132, Sigma) and kept attached to the corresponding carcass for body length determination. Guts were imaged using a stereomicroscope (M205FA, Leica). Gut length was measured and normalized as the gut length/carcass length ratio to account for size variability between larvae.

For cell area analysis, L3 larval guts were dissected and fixed in Bouin’s solution for 2 min and washed in PBST (PBS with 0.5% Triton X-100). After saturation with Normal Goat Serum (G9023, Sigma), mouse anti-Dlg primary antibody (4F3, DSHB) was incubated, at a 1:1000 dilution overnight at 4_°_C. After washes, Alexa Fluor 555 anti-mouse conjugated secondary antibodies (Molecular Probes) were added at 1:200 for 2 h at room temperature. After final washes, samples were mounted in Vectashield Mounting Medium with DAPI (H-1800, Vector Laboratories) and images were acquired as mosaics using the DM6000 (Leica) and analyzed with the Fiji software (Version 1.54r).

### Statistical analysis

Statistical analysis and data visualization were performed using GraphPad Prism 10 software (version 10.6.1). The number of biological replicates (*n*), specific statistical tests, and error bar definitions are detailed in the corresponding figure legends. *P* values are indicated directly on the figures.

### Graphics

Schematics were created with BioRender.

### Declaration of generative AI and AI-assisted technologies in the writing process

During the preparation of this work, the authors used ChatGPT.5 in order to polish the language of the manuscript. After using this tool, the authors reviewed and edited the content as needed and take full responsibility for the content of the publication.

## Supporting information

Suppelemtary materials

## Acknowledgements

The authors would like to thank Berra Erkosar for initial observations; Coralie Drelon for experimental support and Pauline Joncour for bioinformatic support; Naoki Yamanaka for sharing reagents at the early stage of the project; Aurelio Teleman and Gilles Storelli for discussions; the IGFL sequencing and imaging facilities; the PLATIM and Arthro-Tools platforms at the SFR Biosciences Lyon Gerland (UAR3444/US8) for providing Imaging and *Drosophila* facilities; the Bloomington Drosophila Stock Center (BDSC) and Vienna Drosophila Ressource Center (VDRC) for fly stocks and Developmental Studies Hybridoma Bank (DSHB) for antibodies.

## Author contributions

Conceptualization: F.L. and C.I.R.; Methodology: F.L., C.I.R., L.B., S.B. and M.S.; Validation: F.L., C.I.R., L.B. and S.B.; Formal analysis: L.B., C.I.R.; Investigation: L.B., E.G., B.G. and C.I.R.; Resources: C.I.R., L.B., and B.G.; Data curation: L.B. and C.I.R.; Writing - original draft: L.B., C.I.R. and F.L.; Writing - review & editing: L.B., C.I.R. and F.L.; Visualization: L.B. and C.I.R; Supervision: F.L. and C.I.R.; Project administration: C.I.R and F.L..; Funding acquisition: F.L. and C.I.R.

## Supplementary Figures legends

**Figure S1 (related to Figure 1).**
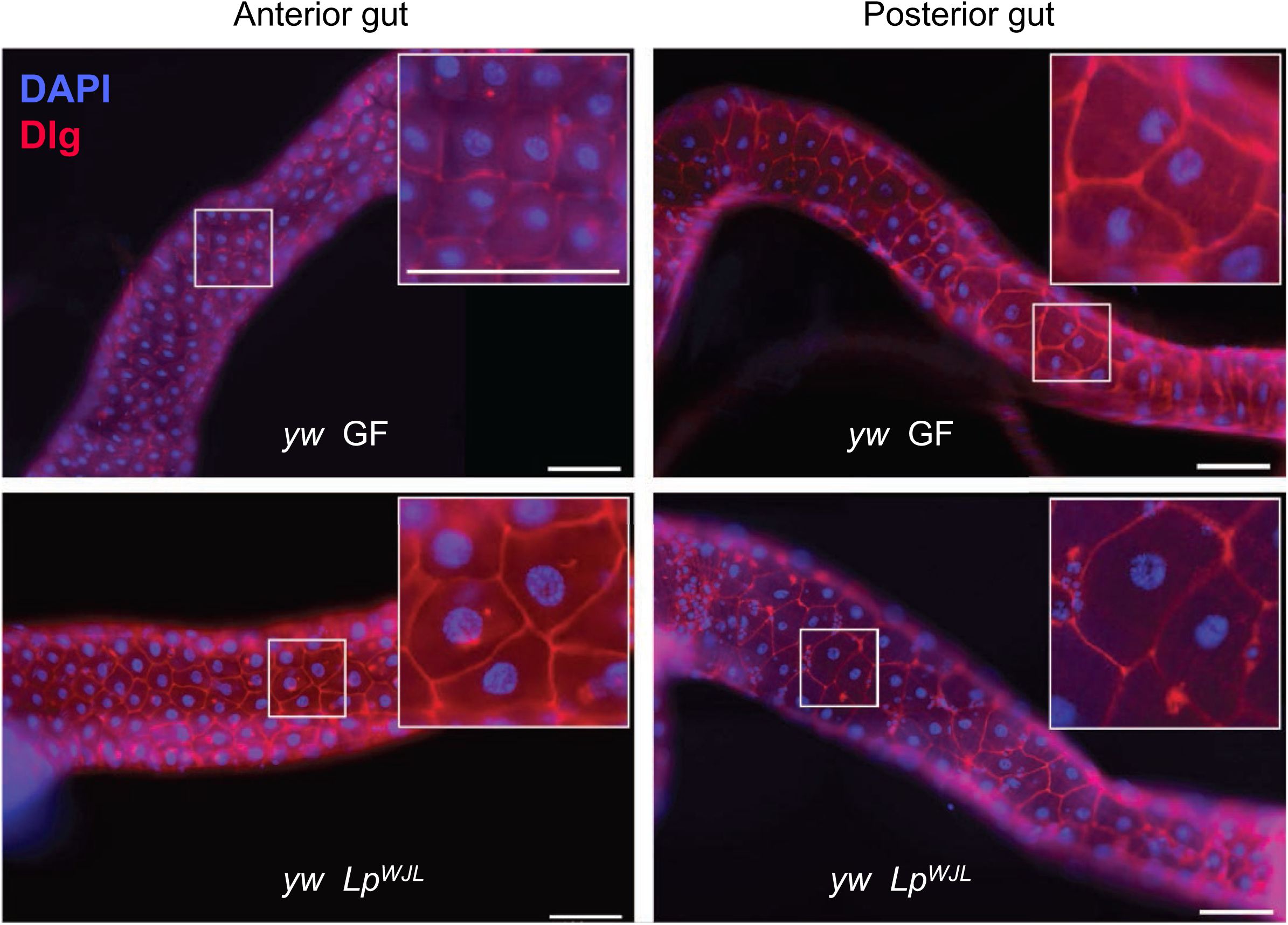
*Lp^WJL^* mediates midgut growth through cell area increase. Representative images of anterior and posterior midguts from size-matched *Lp^WJL^-*associated (D7) and GF (D11) *yw* larvae. Midguts were dissected and stained for Discs large (Dlg, red), a septate junction marker, and DAPI (blue) to mark nuclei. Scale bar represents 100 µm.

**Figure S2 (related to Figure 2).**
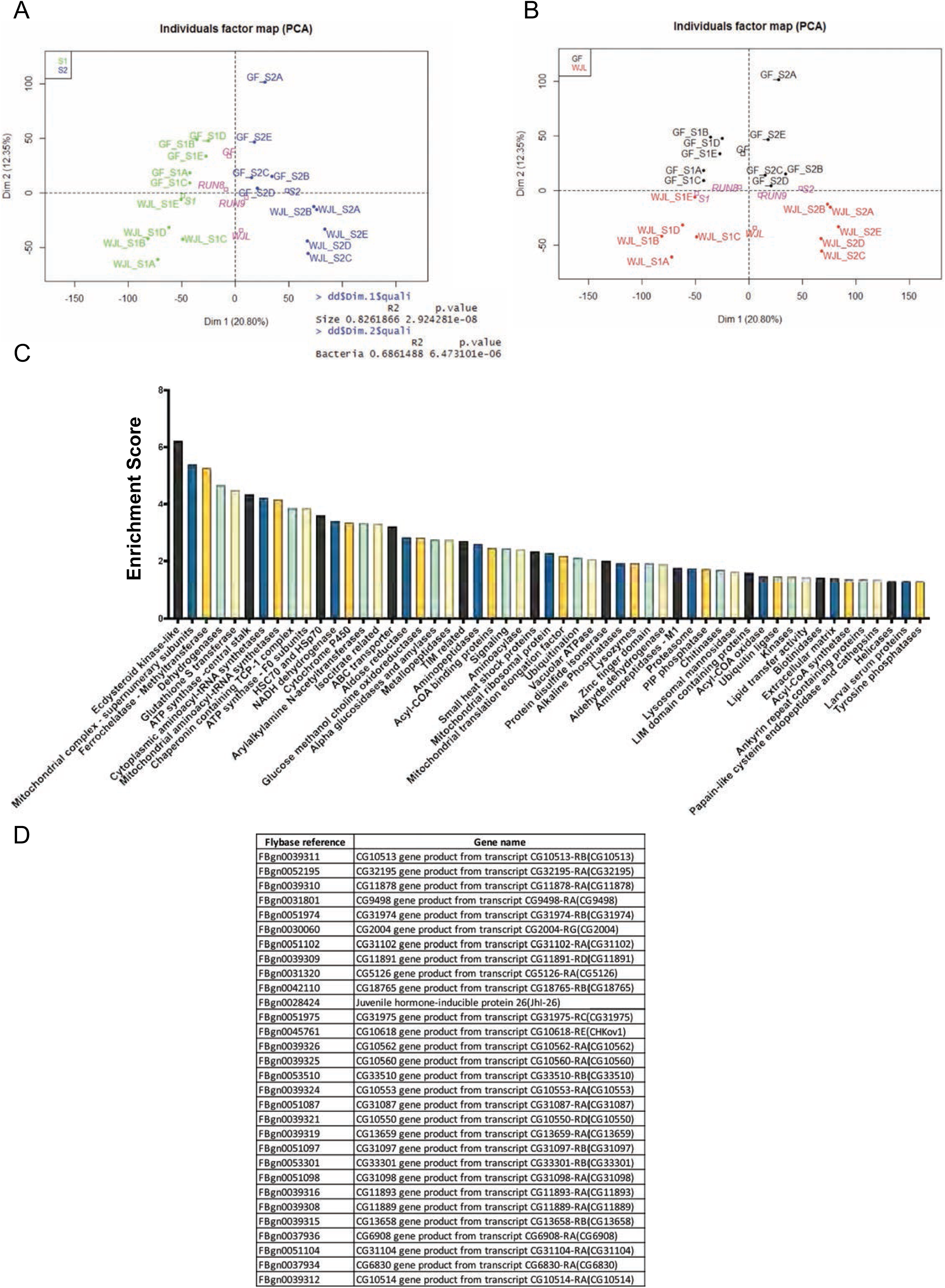
Midgut transcriptomic analysis pinpointes *Lp^WJL^*-triggered EcKL cluster. (A–B) Principal Component Analysis (PCA) plots in (A) show the samples grouped by size, distinguishing Size 1 (green) and Size 2 (blue), with the computed centers for GF and *Lp^WJL^* samples indicated in purple, while the PCA in (B) plots samples based on microbial association, distinguishing *Lp^WJL^*-associated (red) and GF (black) samples, with the computed centers for Size 1 and Size 2 within the GF and *Lp^WJL^* groups shown in purple. (C–D) RNA sequencing results identified 2633 annotated genes that were differentially regulated by *Lp^WJL^* in both sizes; a total of 53 gene clusters (out of 91) were found to be enriched with a score (ES) > 1.3 (*P* < 0.05) (C), and genes belonging to the “EcKL” (Ecdysteroid Kinase like) cluster showed the highest enrichment score.

**Figure S3 (related to Figure 2).**
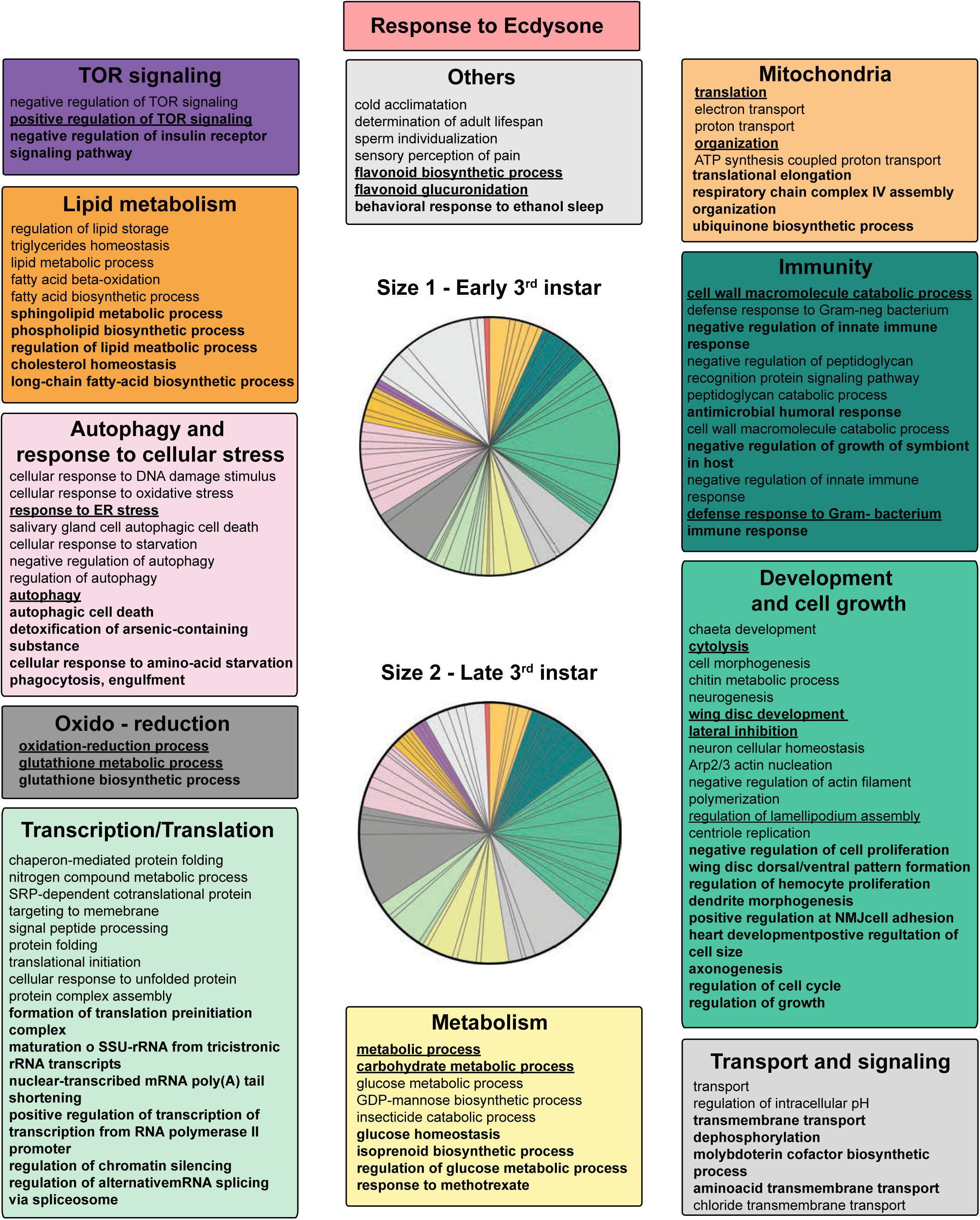
Ecd-responsive biological processes in early L3 and late L3 undernourished larvae. Biological Processes (BP Direct, *P*_adj_ < 5.1ξ 10^-2^) responsive to Ecd are summarized in pie charts for early L3 and late L3 larvae. The pie charts feature boxes highlighting specific BPs: where those found only in early L3 are in normal font, those found only in late L3 are in bold, and those common to both stages are in bold with underlines.

**Figure S4 (related to Figure 2).**
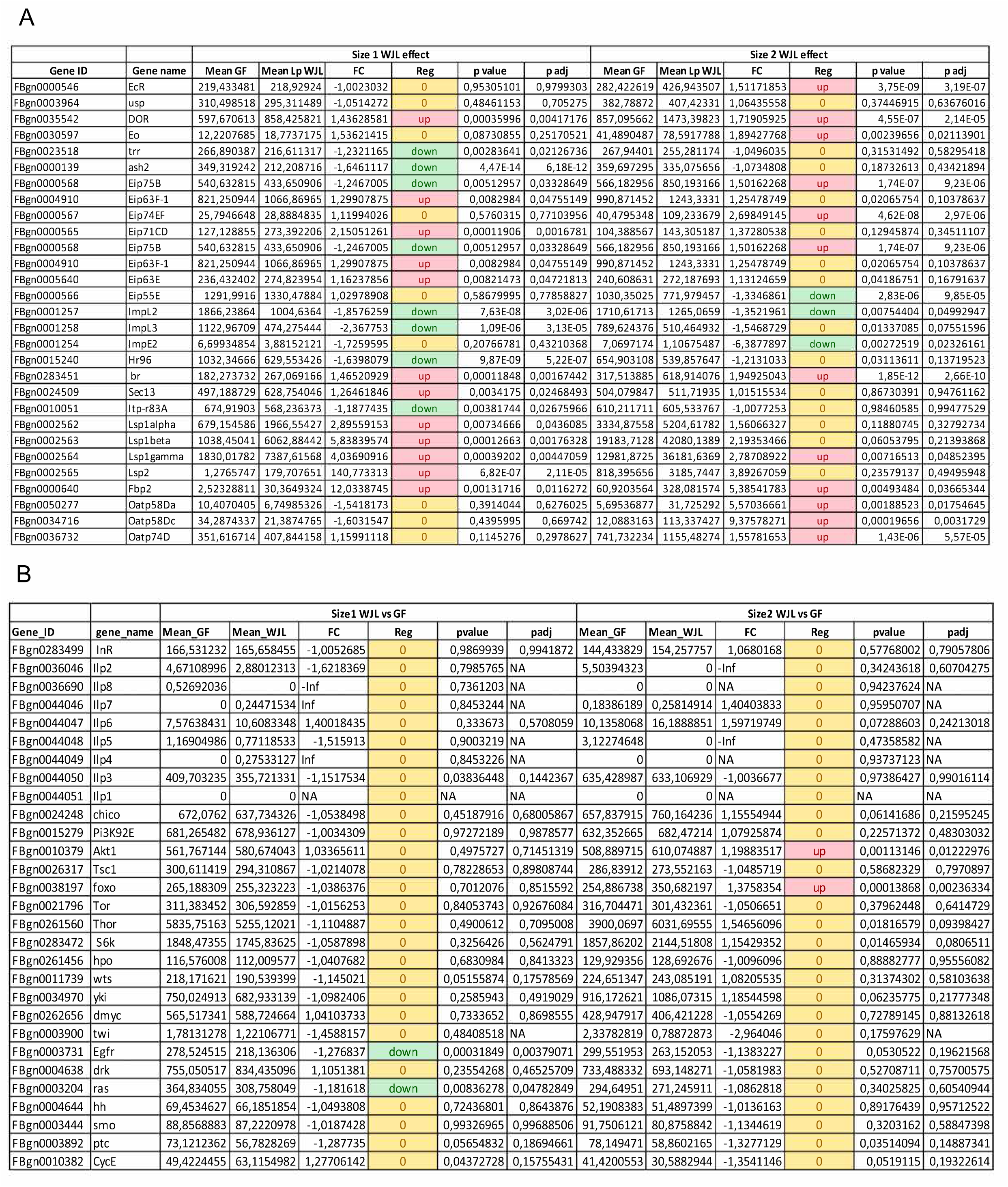
Ecd and tissue growth effector gene expression under *Lp^WJL^* association. Panels (A) and (B) show gene expression data illustrating the differential effect of *Lp^WJL^*association versus GF conditions in L3 larvae. Specifically, the Ecd related genes and the “EcKL” cluster genes were listed in (A), while canonical effectors involved in tissue growth are listed in (B). Mean read counts for both conditions (“Mean GF” and “Mean *Lp^WJL^*”) and the Fold Change (FC) are shown. The mean of the reads counted are presented as “Mean GF” and “Mean *Lp^WJL^*”. FC is indicated by a color code (red for upregulation, yellow for no change, and green for downregulation), with accompanying raw *P* values and adjusted *P* values (*P*_adj_).

**Figure S5 (related to Figure 3).**
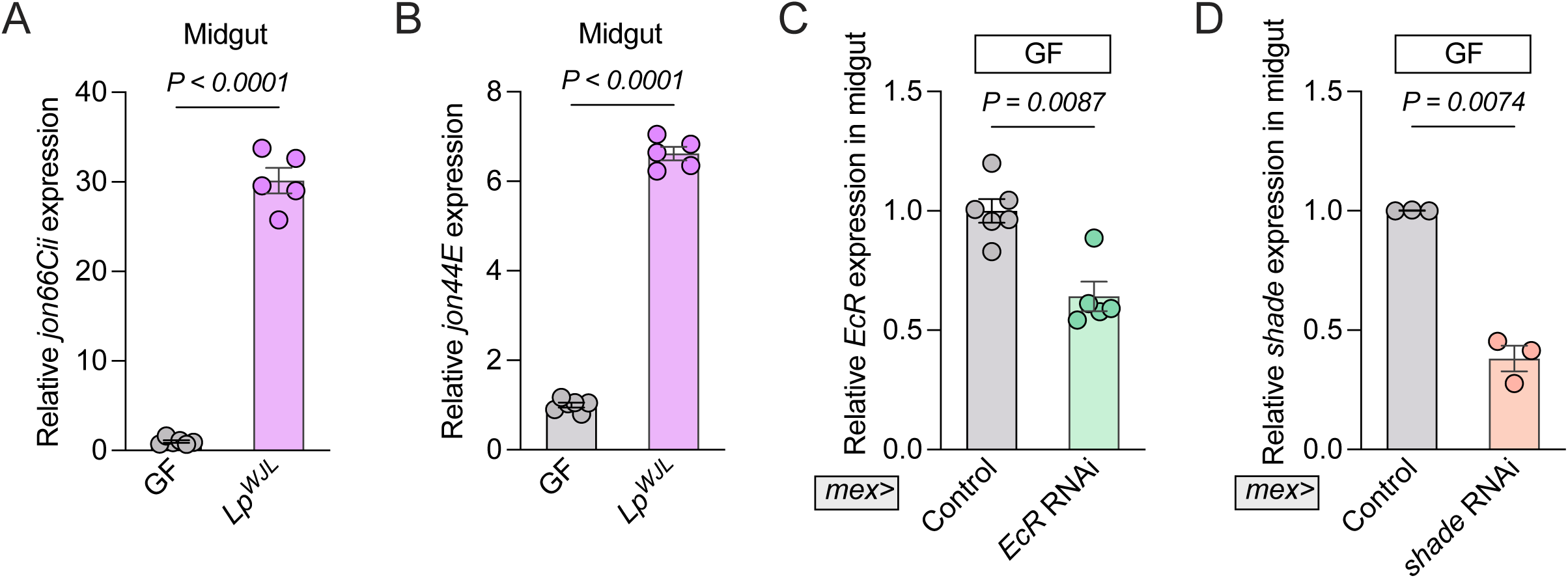
Validation of intestinal peptidases gene expression and knockdown efficiency of Ecdysone effectors. (A–B) The transcript levels of intestinal peptidases *jon66Cii* (A) and *jon44E* (B) were analyzed by qRT-PCR in undernourished GF and *Lp^WJL^*-associated *yw* larvae. n ≥ 5. (C–D) The RNAi efficiency of *EcR* and *shade* in ECs was validated via qRT-PCR in undernourished GF larvae. n ≥ 3. Data are shown as mean ± SEM. Statistical significance was determined using a two-tailed Welch’s *t* test (A, D), a two-tailed unpaired *t* test (B), and a two-tailed Mann-Whitney *U* test (C). *P* values are indicated on the panels.

**Figure S6 (related to Figure 4).**
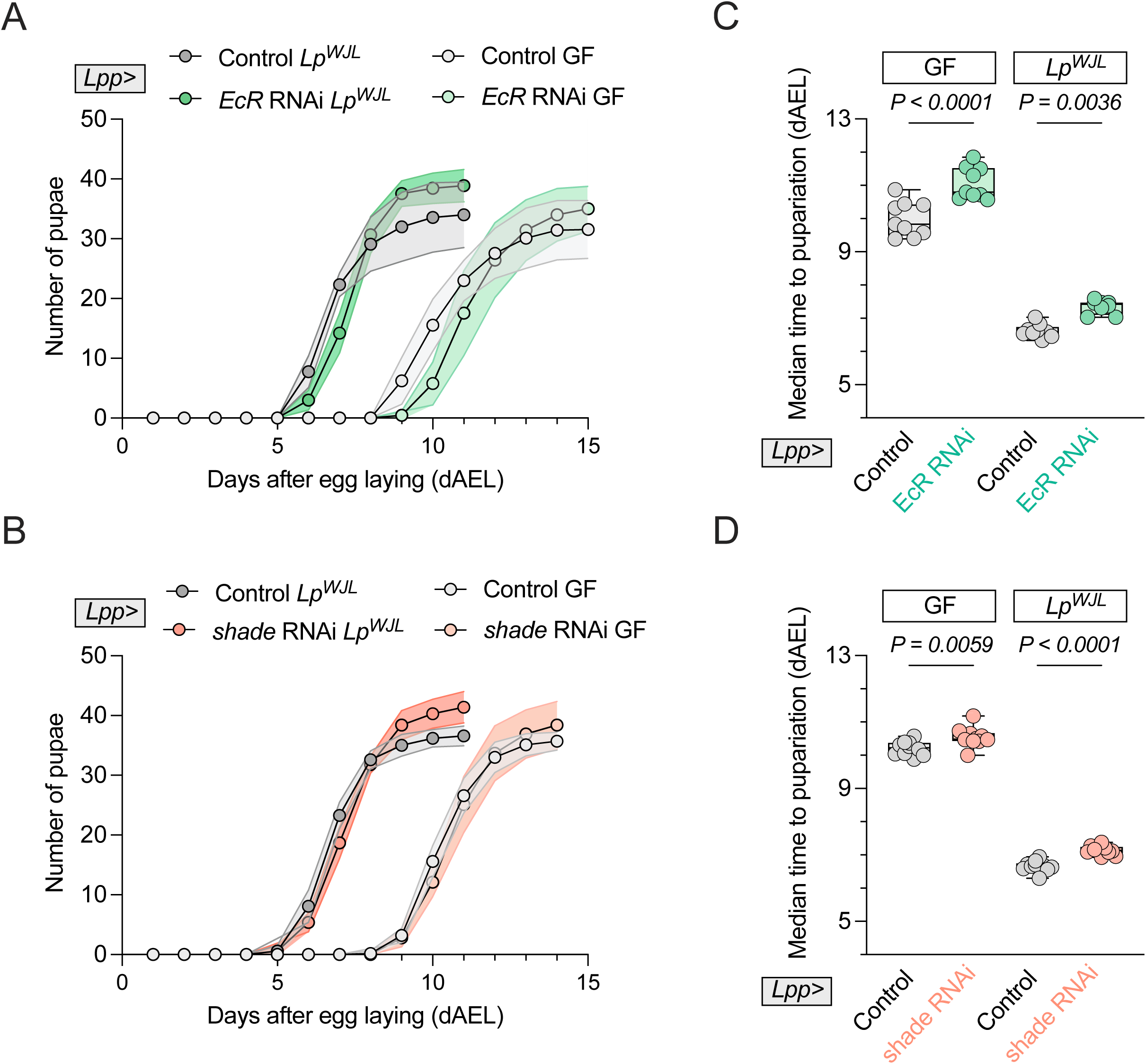
Knowdown of fat body-derived Ecd signaling causes mild developmental delay under malnutrition. (A–D) Under malnutrition, measured developmental timing (A–B) and corresponding median time to pupariation (C–D) are presented in GF and *Lp^WJL^*-associated larvae with *EcR* knockdown in the fat body (*Lpp>EcR* RNAi) (A, C) and *shade* knockdown (*Lpp>shade* RNAi) (B, D). Controls included respective TRiP and KK lines. Statistical significance was determined using a two-way ANOVA with Tukey’s multiple comparisons test. *P* values are indicated on the panels. *n* ≥ 8.

