## Supplementary material for "Ecdysone-mediated intestinal growth contributes to microbiota-driven developmental plasticity under malnutrition": Suppelemtary materials

### SUPPLEMENTARY MATERIALS

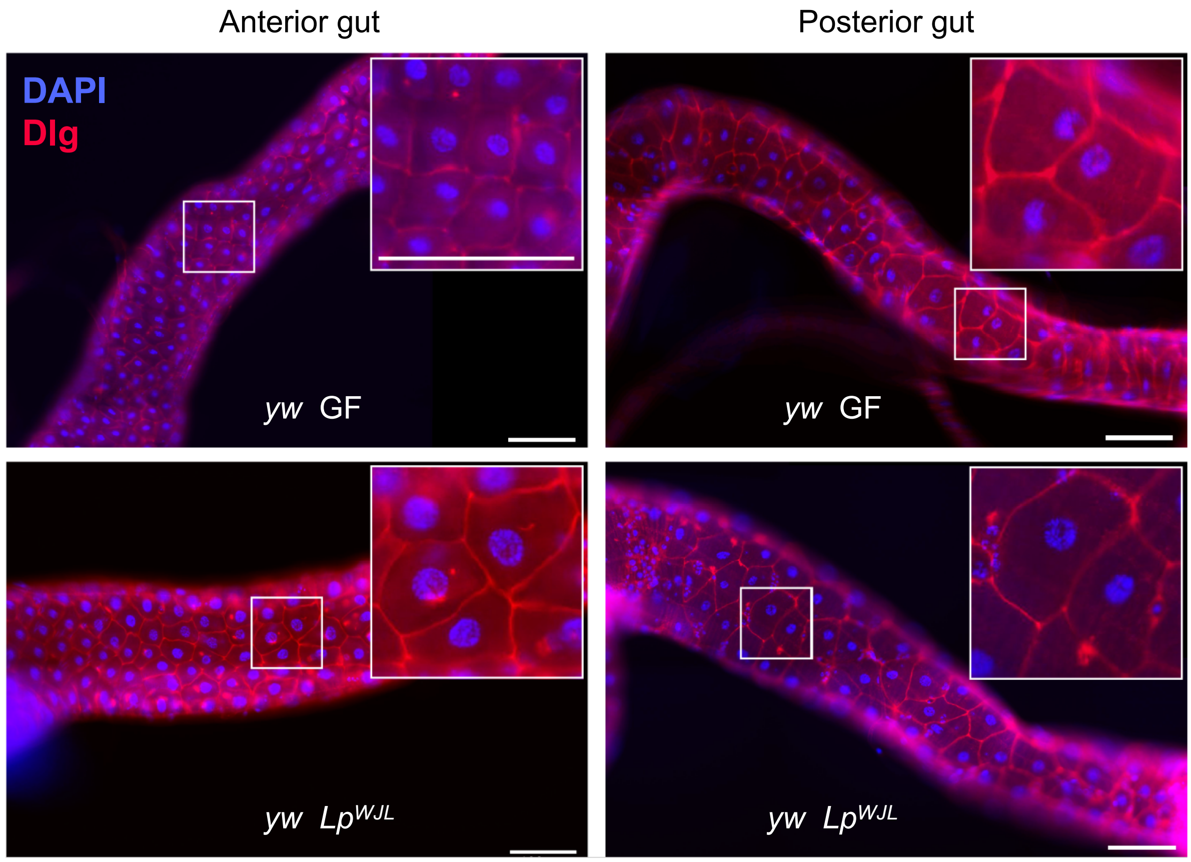


**Figure S1. *Lp*$^{WJL}$ mediates midgut growth through cell area increase.**
Representative images of anterior and posterior midguts from size-matched *Lp*$^{WJL}$-associated (D7) and GF (D11) *yw* larvae. Midguts were dissected and stained for Discs large (Dlg, red), a septate junction marker, and DAPI (blue) to mark nuclei. Scale bar represents 100 µm.

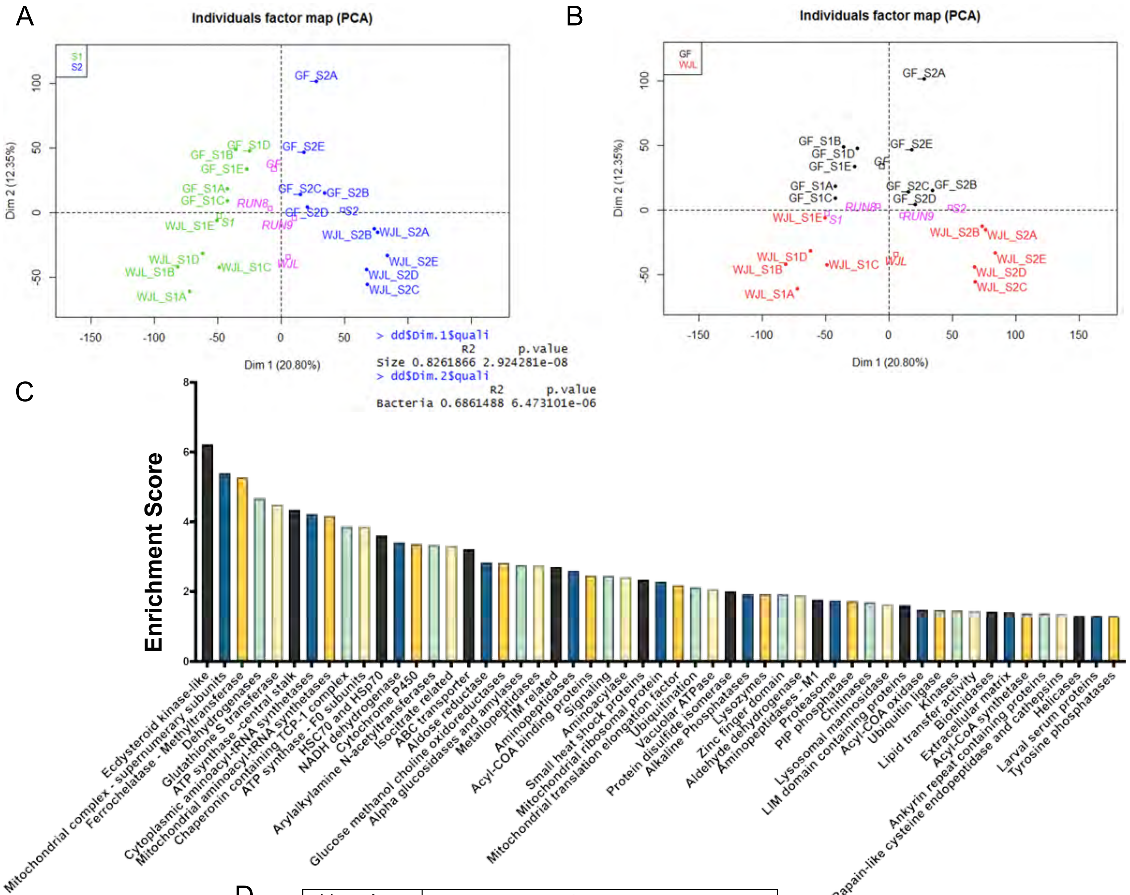


D

| Flybase reference | Gene name |
|---|---|
| FBgn0039311 | CG10513 gene product from transcript CG10513-RB(CG10513) |
| FBgn0052195 | CG32195 gene product from transcript CG32195-RA(CG32195) |
| FBgn0039310 | CG11878 gene product from transcript CG11878-RA(CG11878) |
| FBgn0031801 | CG9498 gene product from transcript CG9498-RA(CG9498) |
| FBgn0051974 | CG31974 gene product from transcript CG31974-RB(CG31974) |
| FBgn0030060 | CG2004 gene product from transcript CG2004-RG(CG2004) |
| FBgn0051102 | CG31102 gene product from transcript CG31102-RA(CG31102) |
| FBgn0039309 | CG11891 gene product from transcript CG11891-RD(CG11891) |
| FBgn0031320 | CG5126 gene product from transcript CG5126-RA(CG5126) |
| FBgn0042110 | CG18765 gene product from transcript CG18765-RB(CG18765) |
| FBgn0028424 | Juvenile hormone-inducible protein 26(JhI-26) |
| FBgn0051975 | CG31975 gene product from transcript CG31975-RC(CG31975) |
| FBgn0045761 | CG10618 gene product from transcript CG10618-RE(CHKov1) |
| FBgn0039326 | CG10562 gene product from transcript CG10562-RA(CG10562) |
| FBgn0039325 | CG10560 gene product from transcript CG10560-RA(CG10560) |
| FBgn0053510 | CG33510 gene product from transcript CG33510-RB(CG33510) |
| FBgn0039324 | CG10553 gene product from transcript CG10553-RA(CG10553) |
| FBgn0051087 | CG31087 gene product from transcript CG31087-RA(CG31087) |
| FBgn0039321 | CG10550 gene product from transcript CG10550-RD(CG10550) |
| FBgn0039319 | CG13659 gene product from transcript CG13659-RA(CG13659) |
| FBgn0051097 | CG31097 gene product from transcript CG31097-RB(CG31097) |
| FBgn0053301 | CG33301 gene product from transcript CG33301-RB(CG33301) |
| FBgn0051098 | CG31098 gene product from transcript CG31098-RA(CG31098) |
| FBgn0039316 | CG11893 gene product from transcript CG11893-RA(CG11893) |
| FBgn0039308 | CG11889 gene product from transcript CG11889-RA(CG11889) |
| FBgn0039315 | CG13658 gene product from transcript CG13658-RB(CG13658) |
| FBgn0037936 | CG6908 gene product from transcript CG6908-RA(CG6908) |
| FBgn0051104 | CG31104 gene product from transcript CG31104-RA(CG31104) |
| FBgn0037934 | CG6830 gene product from transcript CG6830-RA(CG6830) |
| FBgn0039312 | CG10514 gene product from transcript CG10514-RA(CG10514) |

**Figure S2. Midgut transcriptomic analysis pinpointes *Lp$^{WJL}$*-triggered EcKL cluster.**

(A–B) Principal Component Analysis (PCA) plots in (A) show the samples grouped by size, distinguishing Size 1 (green) and Size 2 (blue), with the computed centers for GF and *Lp$^{WJL}$* samples indicated in purple, while the PCA in (B) plots samples based on microbial association, distinguishing *Lp$^{WJL}$*-associated (red) and GF (black) samples, with the computed centers for Size 1 and Size 2 within the GF and *Lp$^{WJL}$* groups shown in purple.

(C–D) RNA sequencing results identified 2633 annotated genes that were differentially regulated by *Lp$^{WJL}$* in both sizes; a total of 53 gene clusters (out of 91) were found to be enriched with a score (ES) > 1.3 ($P < 0.05$) (C), and genes belonging to the "EcKL" (Ecdysteroid Kinase like) cluster showed the highest enrichment score.

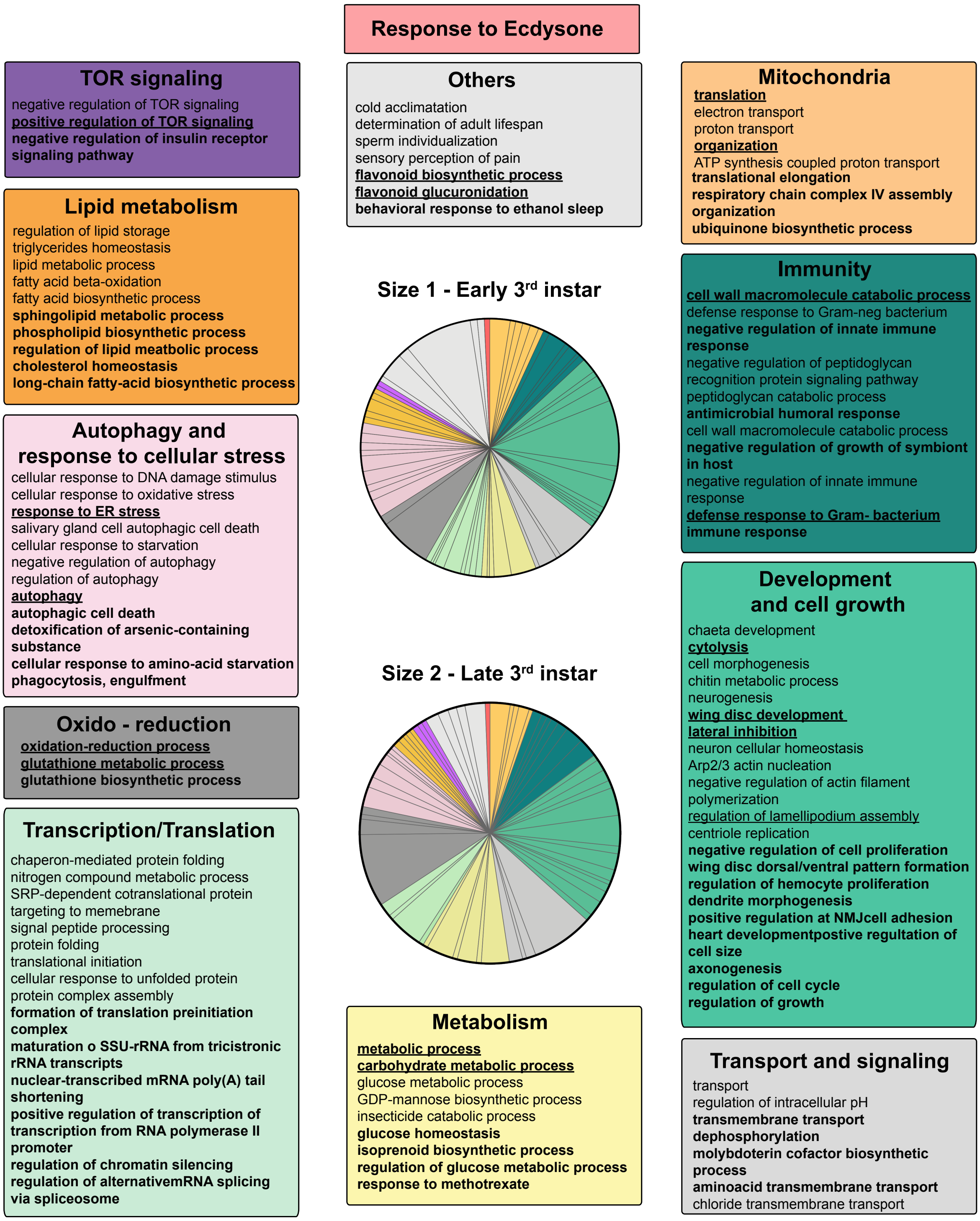


**Figure S3. Ecd-responsive biological processes in early L3 and late L3 undernourished larvae.** Biological Processes (BP Direct, $P_{adj} < 5.1\times 10^{-2}$) responsive to Ecd are summarized in pie charts for early L3 and late L3 larvae. The pie charts feature boxes highlighting specific BPs: where those found only in early L3 are in normal font, those found only in late L3 are in bold, and those common to both stages are in bold with underlines.

A

| | | Size 1 WJL effect | | | | | | Size 2 WJL effect | | | | | |
|---|---|---|---|---|---|---|---|---|---|---|---|---|---|
| Gene ID | Gene name | Mean GF | Mean Lp WJL | FC | Reg | p value | p adj | Mean GF | Mean Lp WJL | FC | Reg | p value | p adj |
| FBgn0000546 | EcR | 219,433481 | 218,92924 | -1,0023032 | 0 | 0,95305101 | 0,9799303 | 282,422619 | 426,943507 | 1,51171853 | up | 3,75E-09 | 3,19E-07 |
| FBgn0003964 | usp | 310,498518 | 295,311489 | -1,0514272 | 0 | 0,48461153 | 0,705275 | 382,78872 | 407,42331 | 1,06435558 | 0 | 0,37446915 | 0,63676016 |
| FBgn0035542 | DOR | 597,670613 | 858,425821 | 1,43628581 | up | 0,00035996 | 0,00417176 | 857,095662 | 1473,39823 | 1,71905925 | up | 4,55E-07 | 2,14E-05 |
| FBgn0030597 | Eo | 12,2207685 | 18,7737175 | 1,53621415 | 0 | 0,08730855 | 0,25170521 | 41,4890487 | 78,5917788 | 1,89427768 | up | 0,00239656 | 0,02113901 |
| FBgn0023518 | trr | 266,890387 | 216,611317 | -1,2321165 | down | 0,00283641 | 0,02126736 | 267,94401 | 255,281174 | -1,0496035 | 0 | 0,31531492 | 0,58295418 |
| FBgn0000139 | ash2 | 349,319242 | 212,208716 | -1,6461117 | down | 4,47E-14 | 6,18E-12 | 359,697295 | 335,075656 | -1,0734808 | 0 | 0,18732613 | 0,43421894 |
| FBgn0000568 | Eip75B | 540,632815 | 433,650906 | -1,2467005 | down | 0,00512957 | 0,03328649 | 566,182956 | 850,193166 | 1,50162268 | up | 1,74E-07 | 9,23E-06 |
| FBgn0004910 | Eip63F-1 | 821,250944 | 1066,86965 | 1,29907875 | up | 0,0082984 | 0,04755149 | 990,871452 | 1243,3331 | 1,25478749 | 0 | 0,02065754 | 0,10378637 |
| FBgn0000567 | Eip74EF | 25,7946648 | 28,8884835 | 1,11994026 | 0 | 0,5760315 | 0,77103956 | 40,4795348 | 109,233679 | 2,69849145 | up | 4,62E-08 | 2,97E-06 |
| FBgn0000565 | Eip71CD | 127,128855 | 273,392206 | 2,15051261 | up | 0,00011906 | 0,0016781 | 104,388567 | 143,305187 | 1,37280538 | 0 | 0,12945874 | 0,34511107 |
| FBgn0000568 | Eip75B | 540,632815 | 433,650906 | -1,2467005 | down | 0,00512957 | 0,03328649 | 566,182956 | 850,193166 | 1,50162268 | up | 1,74E-07 | 9,23E-06 |
| FBgn0004910 | Eip63F-1 | 821,250944 | 1066,86965 | 1,29907875 | up | 0,0082984 | 0,04755149 | 990,871452 | 1243,3331 | 1,25478749 | 0 | 0,02065754 | 0,10378637 |
| FBgn0005640 | Eip63E | 236,432402 | 274,823954 | 1,16237856 | up | 0,00821473 | 0,04721813 | 240,608631 | 272,187693 | 1,13124659 | 0 | 0,04186751 | 0,16791637 |
| FBgn0000566 | Eip55E | 1291,9916 | 1330,47884 | 1,02978908 | 0 | 0,58679995 | 0,77858827 | 1030,35025 | 771,979457 | -1,3346861 | down | 2,83E-06 | 9,85E-05 |
| FBgn0001257 | ImpL2 | 1866,23864 | 1004,6364 | -1,8576259 | down | 7,63E-08 | 3,02E-06 | 1710,61713 | 1265,0659 | -1,3521961 | down | 0,00754404 | 0,04992947 |
| FBgn0001258 | ImpL3 | 1122,96709 | 474,275444 | -2,367753 | down | 1,09E-06 | 3,13E-05 | 789,624376 | 510,464932 | -1,5468729 | 0 | 0,01337085 | 0,07551596 |
| FBgn0001254 | ImpE2 | 6,69934854 | 3,88152121 | -1,7259595 | 0 | 0,20766781 | 0,43210368 | 7,0697174 | 1,10675487 | -6,3877897 | down | 0,00272519 | 0,02326161 |
| FBgn0015240 | Hr96 | 1032,34666 | 629,553426 | -1,6398079 | down | 9,87E-09 | 5,22E-07 | 654,903108 | 539,857647 | -1,2131033 | 0 | 0,03113611 | 0,13719523 |
| FBgn0283451 | br | 182,273732 | 267,069166 | 1,46520929 | up | 0,00011848 | 0,00167442 | 317,513885 | 618,914076 | 1,94925043 | up | 1,85E-12 | 2,66E-10 |
| FBgn0024509 | Sec13 | 497,188729 | 628,754046 | 1,26461846 | up | 0,0034175 | 0,02468493 | 504,079847 | 511,71935 | 1,01515534 | 0 | 0,86730391 | 0,94761162 |
| FBgn0010051 | Itp-r83A | 674,91903 | 568,236373 | -1,1877435 | down | 0,00381744 | 0,02675966 | 610,211711 | 605,533767 | -1,0077253 | 0 | 0,98460585 | 0,99477529 |
| FBgn0002562 | Lsp1alpha | 679,154586 | 1966,55427 | 2,89559153 | up | 0,00734666 | 0,0436085 | 3334,87558 | 5204,61782 | 1,56066327 | 0 | 0,11880745 | 0,32792734 |
| FBgn0002563 | Lsp1beta | 1038,45041 | 6062,88442 | 5,83839574 | up | 0,00012663 | 0,00176328 | 19183,7128 | 42080,1389 | 2,19353466 | 0 | 0,06053795 | 0,21393868 |
| FBgn0002564 | Lsp1gamma | 1830,01782 | 7387,61568 | 4,03690916 | up | 0,00039202 | 0,00447059 | 12981,8725 | 36181,6369 | 2,78708922 | up | 0,00716513 | 0,04852395 |
| FBgn0002565 | Lsp2 | 1,2765747 | 179,707651 | 140,773313 | up | 6,82E-07 | 2,11E-05 | 818,395656 | 3185,7447 | 3,89267059 | 0 | 0,23579137 | 0,49495948 |
| FBgn0000640 | Fbp2 | 2,52328811 | 30,3649324 | 12,0338745 | up | 0,00131716 | 0,0116272 | 60,9203564 | 328,081574 | 5,38541783 | up | 0,00493484 | 0,03665344 |
| FBgn0050277 | Oatp58Da | 10,4070405 | 6,74985326 | -1,5418173 | 0 | 0,3914044 | 0,6276025 | 5,69536877 | 31,725292 | 5,57036661 | up | 0,00188523 | 0,01754645 |
| FBgn0034716 | Oatp58Dc | 34,2874337 | 21,3874765 | -1,6031547 | 0 | 0,4395995 | 0,669742 | 12,0883163 | 113,337427 | 9,37578271 | up | 0,00019656 | 0,0031729 |
| FBgn0036732 | Oatp74D | 351,616714 | 407,844158 | 1,15991118 | 0 | 0,1145276 | 0,2978627 | 741,732234 | 1155,48274 | 1,55781653 | up | 1,43E-06 | 5,57E-05 |

B

| | | Size1 WJL vs GF | | | | | | Size2 WJL vs GF | | | | | |
|---|---|---|---|---|---|---|---|---|---|---|---|---|---|
| Gene_ID | gene_name | Mean_GF | Mean_WJL | FC | Reg | pvalue | padj | Mean_GF | Mean_WJL | FC | Reg | pvalue | padj |
| FBgn0283499 | InR | 166,531232 | 165,658455 | -1,0052685 | 0 | 0,9869939 | 0,9941872 | 144,433829 | 154,257757 | 1,0680168 | 0 | 0,57768002 | 0,79057806 |
| FBgn0036046 | Ilp2 | 4,67108996 | 2,88012313 | -1,6218369 | 0 | 0,7985765 | NA | 5,50394323 | 0 | -Inf | 0 | 0,34243618 | 0,60704275 |
| FBgn0036690 | Ilp8 | 0,52692036 | 0 | -Inf | 0 | 0,7361203 | NA | 0 | 0 | NA | 0 | 0,94237624 | NA |
| FBgn0044046 | Ilp7 | 0 | 0,24471534 | Inf | 0 | 0,8453244 | NA | 0,18386189 | 0,25814914 | 1,40403833 | 0 | 0,95950707 | NA |
| FBgn0044047 | Ilp6 | 7,57638431 | 10,6083348 | 1,40018435 | 0 | 0,333673 | 0,5708059 | 10,1358068 | 16,1888851 | 1,59719749 | 0 | 0,07288603 | 0,24213018 |
| FBgn0044048 | Ilp5 | 1,16904986 | 0,77118533 | -1,515913 | 0 | 0,9003219 | NA | 3,12274648 | 0 | -Inf | 0 | 0,47358582 | NA |
| FBgn0044049 | Ilp4 | 0 | 0,27533127 | Inf | 0 | 0,8453226 | NA | 0 | 0 | NA | 0 | 0,93737123 | NA |
| FBgn0044050 | Ilp3 | 409,703235 | 355,721331 | -1,1517534 | 0 | 0,03836448 | 0,1442367 | 635,428987 | 633,106929 | -1,0036677 | 0 | 0,97386427 | 0,99016114 |
| FBgn0044051 | Ilp1 | 0 | 0 | NA | 0 | NA | NA | 0 | 0 | NA | 0 | NA | NA |
| FBgn0024248 | chico | 672,0762 | 637,734326 | -1,0538498 | 0 | 0,45187916 | 0,68005867 | 657,837915 | 760,164236 | 1,15554944 | 0 | 0,06141686 | 0,21595245 |
| FBgn0015279 | Pi3K92E | 681,265482 | 678,936127 | -1,0034309 | 0 | 0,97272189 | 0,9878577 | 632,352665 | 682,47214 | 1,07925874 | 0 | 0,22571372 | 0,48303032 |
| FBgn0010379 | Akt1 | 561,767144 | 580,674043 | 1,03365611 | 0 | 0,4975727 | 0,71451319 | 508,889715 | 610,074887 | 1,19883517 | up | 0,00113146 | 0,01222976 |
| FBgn0026317 | Tsc1 | 300,611419 | 294,310867 | -1,0214078 | 0 | 0,78228653 | 0,89808744 | 286,83912 | 273,552163 | -1,0485719 | 0 | 0,58682329 | 0,7970897 |
| FBgn0038197 | foxo | 265,188309 | 255,323223 | -1,0386376 | 0 | 0,7012076 | 0,8515592 | 254,886738 | 350,682197 | 1,3758354 | up | 0,00013868 | 0,00236334 |
| FBgn0021796 | Tor | 311,383452 | 306,592859 | -1,0156253 | 0 | 0,84053743 | 0,92676084 | 316,704471 | 301,432361 | -1,0506651 | 0 | 0,37962448 | 0,6414729 |
| FBgn0261560 | Thor | 5835,75163 | 5255,12021 | -1,1104887 | 0 | 0,4900612 | 0,7095008 | 3900,0697 | 6031,69555 | 1,54656096 | 0 | 0,01816579 | 0,09398427 |
| FBgn0283472 | S6k | 1848,47355 | 1745,83625 | -1,0587898 | 0 | 0,3256426 | 0,5624791 | 1857,86202 | 2144,51808 | 1,15429352 | 0 | 0,01465934 | 0,0806511 |
| FBgn0261456 | hpo | 116,576008 | 112,009577 | -1,0407682 | 0 | 0,6830984 | 0,8413323 | 129,929356 | 128,692676 | -1,0096096 | 0 | 0,88882777 | 0,95556082 |
| FBgn0011739 | wts | 218,171621 | 190,539399 | -1,145021 | 0 | 0,05155874 | 0,17578569 | 224,651347 | 243,085191 | 1,08205535 | 0 | 0,31374302 | 0,58103638 |
| FBgn0034970 | yki | 750,024913 | 682,933139 | -1,0982406 | 0 | 0,2585943 | 0,4919029 | 916,172621 | 1086,07315 | 1,18544598 | 0 | 0,06235775 | 0,21777348 |
| FBgn0262656 | dmyc | 565,517341 | 588,724664 | 1,04103733 | 0 | 0,7333652 | 0,8698555 | 428,947917 | 406,421228 | -1,0554269 | 0 | 0,72789145 | 0,88132618 |
| FBgn0003900 | twi | 1,78131278 | 1,22106771 | -1,4588157 | 0 | 0,48408518 | NA | 2,33782819 | 0,78872873 | -2,964046 | 0 | 0,17597629 | NA |
| FBgn0003731 | Egfr | 278,524515 | 218,136306 | -1,276837 | down | 0,00031849 | 0,00379071 | 299,551953 | 263,152053 | -1,1383227 | 0 | 0,0530522 | 0,19621568 |
| FBgn0004638 | drk | 755,050517 | 834,435096 | 1,1051381 | 0 | 0,23554268 | 0,46525709 | 733,488332 | 693,148271 | -1,0581983 | 0 | 0,52708711 | 0,75700575 |
| FBgn0003204 | ras | 364,834055 | 308,758049 | -1,181618 | down | 0,00836278 | 0,04782849 | 294,64951 | 271,245911 | -1,0862818 | 0 | 0,34025825 | 0,60540944 |
| FBgn0004644 | hh | 69,4534627 | 66,1851854 | -1,0493808 | 0 | 0,72436801 | 0,8643876 | 52,1908383 | 51,4897399 | -1,0136163 | 0 | 0,89176439 | 0,95712522 |
| FBgn0003444 | smo | 88,8568883 | 87,2220978 | -1,0187428 | 0 | 0,99326965 | 0,99688506 | 91,7506121 | 80,8758842 | -1,1344619 | 0 | 0,3203162 | 0,58847398 |
| FBgn0003892 | ptc | 73,1212362 | 56,7828269 | -1,287735 | 0 | 0,05654832 | 0,18694661 | 78,149471 | 58,8602165 | -1,3277129 | 0 | 0,03514094 | 0,14887341 |
| FBgn0010382 | CycE | 49,4224455 | 63,1154982 | 1,27706142 | 0 | 0,04372728 | 0,15755431 | 41,4200553 | 30,5882944 | -1,3541146 | 0 | 0,0519115 | 0,19322614 |

**Figure S4. Ecd and tissue growth effector gene expression under *Lp*$^{WJL}$ association.**

Panels (A) and (B) show gene expression data illustrating the differential effect of *Lp*$^{WJL}$ association versus GF conditions in L3 larvae. Specifically, the Ecd related genes and the "EcKL" cluster genes were listed in (A), while canonical effectors involved in tissue growth are listed in (B). Mean read counts for both conditions ("Mean GF" and "Mean *Lp*$^{WJL}$") and the Fold Change (FC) are shown. The mean of the reads counted are presented as "Mean GF" and "Mean *Lp*$^{WJL}$". FC is indicated by a color code (red for upregulation, yellow for no change, and green for downregulation), with accompanying raw *P* values and adjusted *P* values ($P_{adj}$).

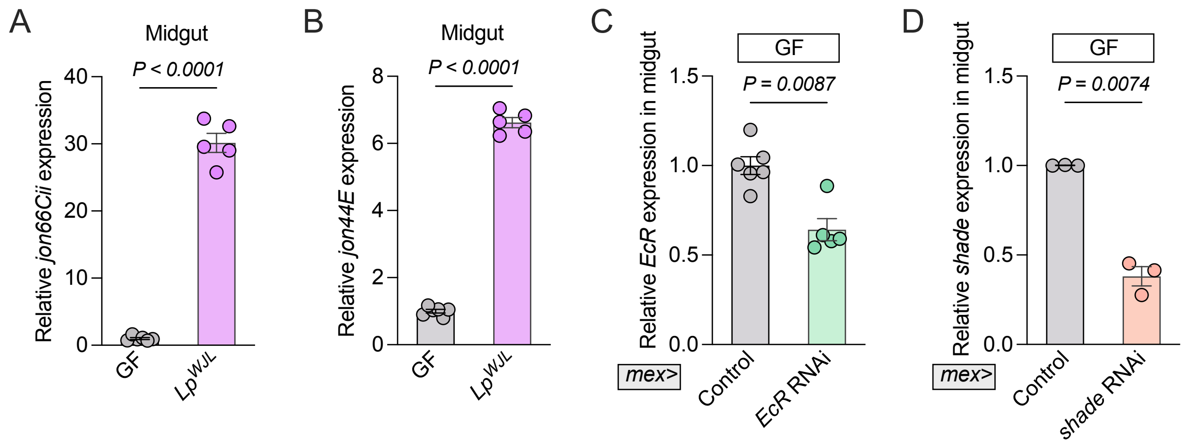


**Figure S5. Validation of intestinal peptidases gene expression and knockdown efficiency of Ecdysone effectors.**

(A–B) The transcript levels of intestinal peptidases *jon66Cii* (A) and *jon44E* (B) were analyzed by qRT-PCR in undernourished GF and $Lp^{WJL}$-associated *yw* larvae. n ≥ 5.

(C–D) The RNAi efficiency of *EcR* and *shade* in ECs was validated via qRT-PCR in undernourished GF larvae. n ≥ 3.

Data are shown as mean ± SEM. Statistical significance was determined using a two-tailed Welch's *t* test (A, D), a two-tailed unpaired *t* test (B), and a two-tailed Mann-Whitney *U* test (C). *P* values are indicated on the panels.

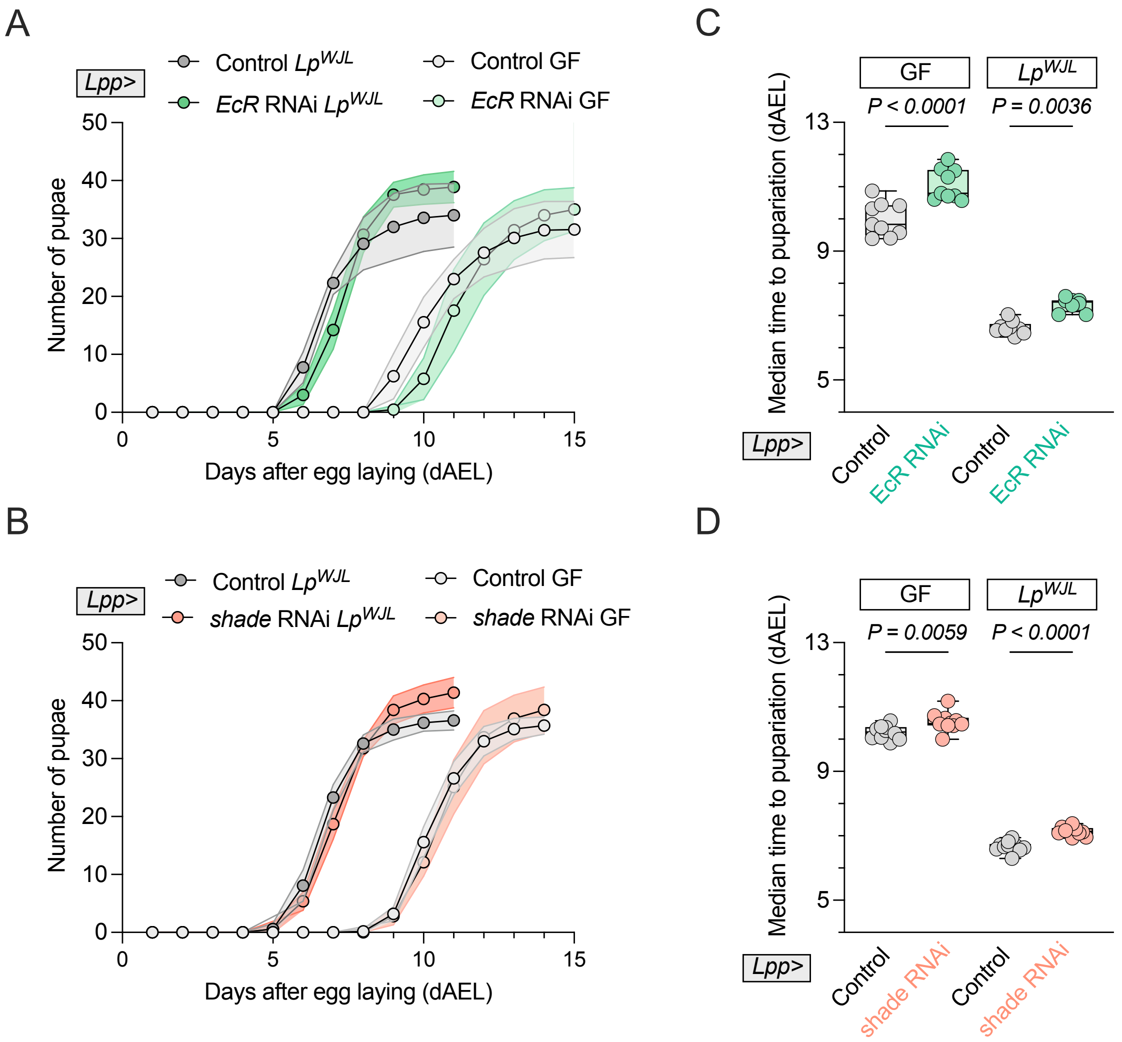


**Figure S6. Knowdown of fat body-derived Ecd signaling causes mild developmental delay under malnutrition.**

(A–D) Under malnutrition, measured developmental timing (A–B) and corresponding median time to pupariation (C–D) are presented in GF and $Lp^{WJL}$-associated larvae with *EcR* knockdown in the fat body (*Lpp>EcR* RNAi) (A, C) and *shade* knockdown (*Lpp>shade* RNAi) (B, D). Controls included respective TRiP and KK lines. Statistical significance was determined using a two-way ANOVA with Tukey's multiple comparisons test. *P* values are indicated on the panels. $n \geq 8$.
